# Dynamics of Meiotic Sex Chromosome Inactivation and Pachytene Activation in Mice Spermatogenesis

**DOI:** 10.1101/665372

**Authors:** Ábel Vértesy, Javier Frias-Aldeguer, Zeliha Sahin, Nicolas Rivron, Alexander van Oudenaarden, Niels Geijsen

## Abstract

During germ cell development, cells undergo a drastic switch from mitosis to meiosis to form haploid germ cells. Sequencing and computational technologies now allow studying development at the single-cell level. Here we developed a multiplexed trajectory reconstruction to create a high-resolution developmental map of spermatogonia and prophase-I spermatocytes from testes of a Dazl-GFP reporter mouse. We identified three main transitions in the meiotic prophase-I: meiotic entry, the meiotic sex chromosome inactivation (MSCI), and concomitant pachytene activation. We validated the key features of these transitions *in vivo* using single molecule FISH. Focusing on MSCI, we found that 34% of sex chromosomal genes are induced shortly before MSCI, that silencing time is diverse and correlates with specific gene functions. These highlight a previously underappreciated level of regulation of MSCI. Finally, we found that spermatozoal genes in pachytene are activated in a temporal pattern reflecting the future anatomic and functional order of the sperm cell. Altogether we highlighted how precise and sequential changes in gene expression regulate cellular states in meiotic prophase-I.

## Introduction

Spermatogenesis is a complex and continuous differentiation process from the spermatogonia to the spermatocyte stages that resolves a switch from mitotic to meiotic divisions to generate haploid sperm cells. This process is accompanied by major morphological, cellular and transcriptional changes, which are not well understood transcriptome-wide nor temporally resolved in a fine manner. Single-cell analysis provided an overall description of spermatogenesis (Chen *et al*., 2018; Jung *et al*., 2018; Lukassen *et al*., 2018) including the analysis of Sertoli cell biology (Green *et al*., 2018), the comparison of human and mouse spermatogenesis (Hermann *et al*., 2018) and the analysis of post-meiotic sex chromosome dynamics (Ernst *et al*., 2019). Here, we developed a multiplex computational approach to finely resolve developmental trajectories of single *Dazl+* cells over the time of mitotic to meiosis transition.

A male specific feature of meiosis is the incomplete pairing of the X and Y chromosomes during meiosis (termed asynapsis). While in female cells, all chromosomes can form homologous pairs and recombine, in males, the X and Y chromosomes only share a small homologous part, the pseudoautosomal region (PAR). The remaining non-homologous regions stay unpaired, acquire repressive chromatin marks, and eventually form the sex-body within the nucleus (Khalil, Boyar and Driscoll, 2004; van der Heijden *et al*., 2007). This silencing of the X and Y chromosome during male meiosis is called meiotic sex chromosome inactivation (MSCI) (Henderson, 1963; Turner, 2015). MSCI is thought as a special case of meiotic silencing of unsynapsed chromatin (Shiu *et al*., 2001) and is an essential step of male meiosis (Royo *et al*., 2015). While MSCI’s post transcriptional regulation, such as H2AX phosphorylation (Mahadevaiah *et al*., 2001) and ubiquitination (Baarends *et al*., 2005) are well described, the transcriptional control of these regulators and the dynamics of gene expression silencing are still to be identified.

Another feature of male meiosis is a massive wave of gene expression at the pachytene stage (Fallahi *et al*., 2010), when mid-meiosis cells express a remarkable array of post-meiotic genes involved in spermiogenesis (da Cruz *et al*., 2016). The dynamics of the activation of this transcriptional program, and whether sex chromosomal genes can escape MSCI are open questions in the field, which we analyzed in detail.

We thus set out to draw a temporally resolved, transcriptome-wide gene expression atlas from meiotic commitment until prophase-I, in order to unravel transcriptional changes that 1880 single cells, multiplex reconstructed their temporal order *in silico*, which we validated *via* microscopy. This allowed us to describe the precise sequence of transcriptome-wide changes upon MSCI and pachytene activation.

## Results

### 1. Single-cell transcriptomics of *Dazl+* germ cells identifies stem cells along with 4 stages of prophase-I

*Dazl* is expressed from spermatogonial to early pachytene stage (Kim, Cooke and Rhee, 2012). We thus used *Dazl-GFP* reporter mice (Chen *et al*., 2014) to isolate cells during meiotic entry and prophase from testes of adult mice (2-4 month) (Fig. 1A). We applied single-cell mRNA sequencing using the SORT-seq protocol (Muraro *et al*., 2016), and sequenced 1880 single cells from 3 mice. After trimming and sequence alignment to the mm10 reference transcriptome (Online Methods), we filtered low quality cells with fewer than 3000 unique (UMI corrected) transcripts and selected 1274 high-quality germ cells. These were normalized to the median transcript count (20488 per cell), allowing us to compare the relative expression across cells (Fig. S1A, Online Methods).

**Fig. 1:**
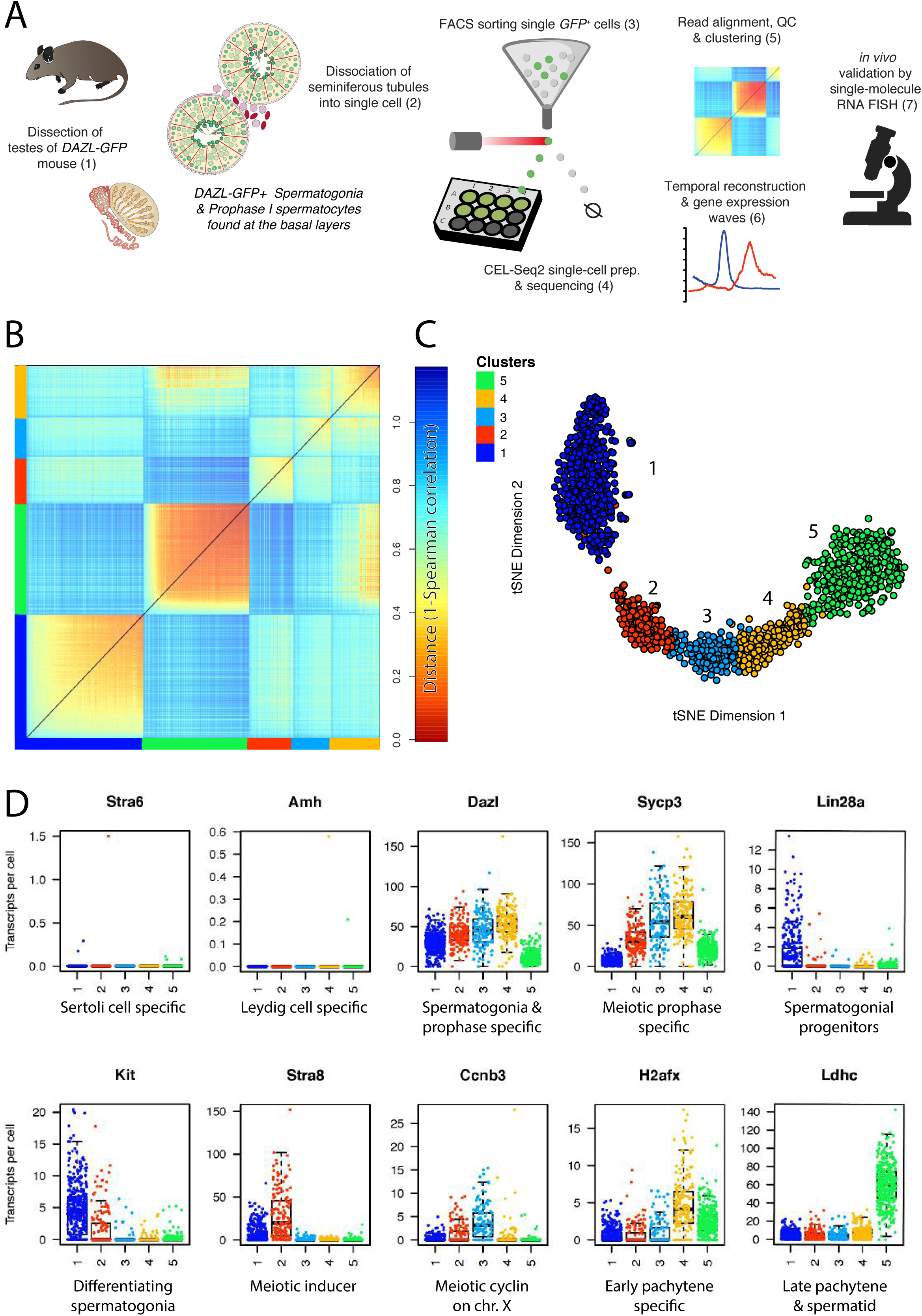
Single cell mRNA sequencing identifies five subtypes of *Dazl*-GFP+ germ cells, which correspond to subsequent stages of prophase-I. a.) Experimental and analytic workflow. b.) Transcriptome-wide k-medoids clustering identifies 5 clusters among 1274 high quality Dazl-GFP+ cells. Spearman correlation coefficients show homogenous and distinct clusters (cl. 1 & 5), while the three other stages are more heterogeneous. c.) Cell-to-cell differences visualized on t-SNE map. Stages along the curve from left (1) to right (5) represent consecutive stages of prophase-I. Colors and numbers indicate clusters (as determined in panel B) and represent spermatogonial (1), (pre)leptotene I (2), zygotene (3), and pachytene (4,5) stages. Each dot represents a single cell. d.) Expression of the selected marker genes. Negative and positive controls: No sorted cells express the Sertoli cell specific retinol binding protein 4 receptor (*Stra6*) or the Leydig cell specific anti-Mullerian hormone (*Amh*), but all cells express the sorting marker Dazl-GFP mRNA, validating sorting approach. Synaptonemal Complex Protein 3 (*Sycp3*) is a core component of the synaptonemal complex connecting homologous chromosomes in meiotic prophase. **Cl. 1** cells express spermatogonial markers *Lin28a* (*progenitors*) or *Kit* (differentiating spermatogonia). **Cl. 2** cells uniquely express the preleptotene marker *Stra8*, a key inducer of meiosis. **Cl. 3** cells the meiosis specific Cyclin B3 (*Ccnb3*) gene, which is located on the X chromosome, marking the last stage where sex chromosomes are expressed (zygotene). **Cl. 4** cells upregulate H2A histone family member X (*H2afx*), the histone variant associated with meiotic sex chromosome inactivation in early pachytene. **Cl. 5** cells specifically express many genes related to the function of spermatids. e.g.: lactate dehydrogenase C (*Ldhc*), testis-specific enzyme in anaerobic glycolysis, producing energy in motile sperm.

To determine whether transcriptional changes are continuous or rather separated into discrete steps, we classified cells by transcriptome-wide, unsupervised k-medoids clustering on Spearman correlation coefficients. Cells were clustered in 5 groups (Fig. 1 B, C) based on all genes with at least 10 transcripts in minimum 50 cells. Importantly, we did not predefine the number of expected stages, but estimated it based on the within-cluster-dispersion of data points (Grün *et al*., 2016). The established clusters are also supported by saturating gap statistics, a positive cluster silhouette. High Jaccard indices across bootstrapping rounds indicate that the clusters are reproducible (Fig. S1 C-E, Online Methods). These clusters are not influenced by animal-to-animal variation, batch variation nor library complexity (Fig. S2). After we ruled out batch effects, we excluded contamination from somatic cells based on a lack of expression of Sertoli or Leydig cell specific markers *Stra6* and *Amh* (Fig. 1D) (O’Shaughnessy, Morris and Baker, 2008; Griswold, 2016). On the contrary, cells showed a high expression of *Dazl*, and of the meiotic prophase marker *Sycp3* (Fig. 1D).

Cells forming cluster 1 express markers of differentiating spermatogonia, like *Dmrt1* (Matson *et al*., 2010), *Kit* (Yoshinaga *et al*., 1991), and earlier progenitor markers, as *Lin28a* (Chakraborty *et al*., 2014) (Fig. 1 D). Cells forming the cluster 2 uniquely express *Stra8*, the key gene marking meiotic entry (Anderson *et al*., 2008). Cells forming cluster 3 specifically upregulate the zygotene specific Cyclin B3 (*Ccnb3*) (Nguyen *et al*., 2002), which is located on the X chromosome, and marks the last stage with sex chromosomal expression (zygotene I). Cells forming cluster 4 upregulated H2A histone family member X (*H2afx*), the histone variant associated with meiotic sex chromosome inactivation (MSCI) in early pachytene. Finally, cells forming cluster 5 specifically express the pachytene marker lactate dehydrogenase C (*Ldhc*), a testis-specific enzyme in anaerobic glycolysis (Goldberg *et al*., 2010). Taken together, we concluded that *Dazl+* cells represent 5 transcriptionally distinct subpopulations with clear transcriptional signatures.

### 2. Identified clusters correspond to subsequent stages of prophase-I

To better characterize the established clusters, we compared their gene expression by differential gene expression analysis (Table S1-S2, Data S2). Cluster 1 on the left edge of the t-SNE map (Fig 1B) encloses ∼33% of the cells (Fig. S2A). They are uniquely expressing *Dmrt1* and other markers of mitotically dividing cells including *Ccnd1, Cenpa, Mcm2*, *Ccdc6, Gsr, Ccdc6 and Crabp1 (*Data S2, Table S1). While most spermatogonial cells are *Kit*+, we noticed that some markers of the undifferentiated state, including *Uchl1* or *Lin28a* (Luo, Megee and Dobrinski, 2009; Griswold, 2016) are partially co-expressed with Kit+ cells (Data S1).

On the right side of the t-SNE map (Fig 1B), 36% of the cells form Cluster 5 (Fig. S2A). These cells are characterized by a high expression of pachytene markers such as the *Adam* genes, *Ldha, Ldhc, Crisp2, Ybx* genes, *Meig1* and *Piwil1* (Jan *et al*., 2017) (Data S2), consistent with early expression of post-meiotic genes (da Cruz *et al*., 2016).

These two opposite clusters (1 and 5) are defined by many uniquely expressed genes within our dataset (Table S1-S2). Between these, we found 3 intermediate clusters with fewer specific genes. These clusters therefore are largely determined by unique combinations of up- and downregulated genes. For example, we found that Cluster 2 uniquely expresses *Stra8* and *Prdm9* suggestive of differentiated spermatogonial or preleptotene stage (Sun *et al*., 2015). At the same time, they also started to upregulate the meiotic *Sycp* genes (Margolin *et al*., 2014).

The meiotic entry is accompanied by a global transcriptional silencing (Monesi, 1965; Ernst *et al*., 2019). Indeed, we observed that the meiotic entry (Cluster 2) corresponds to a transient drop in raw transcript counts in clusters 2-4 (Fig. 2D), which is not explained by differences in sequencing depth (Fig. S1F). Despite this drastic silencing, each cluster (1 & 2) showed the upregulation of a specific subset of genes, as compared to the earlier stage (Table S2).

**Fig. 2:**
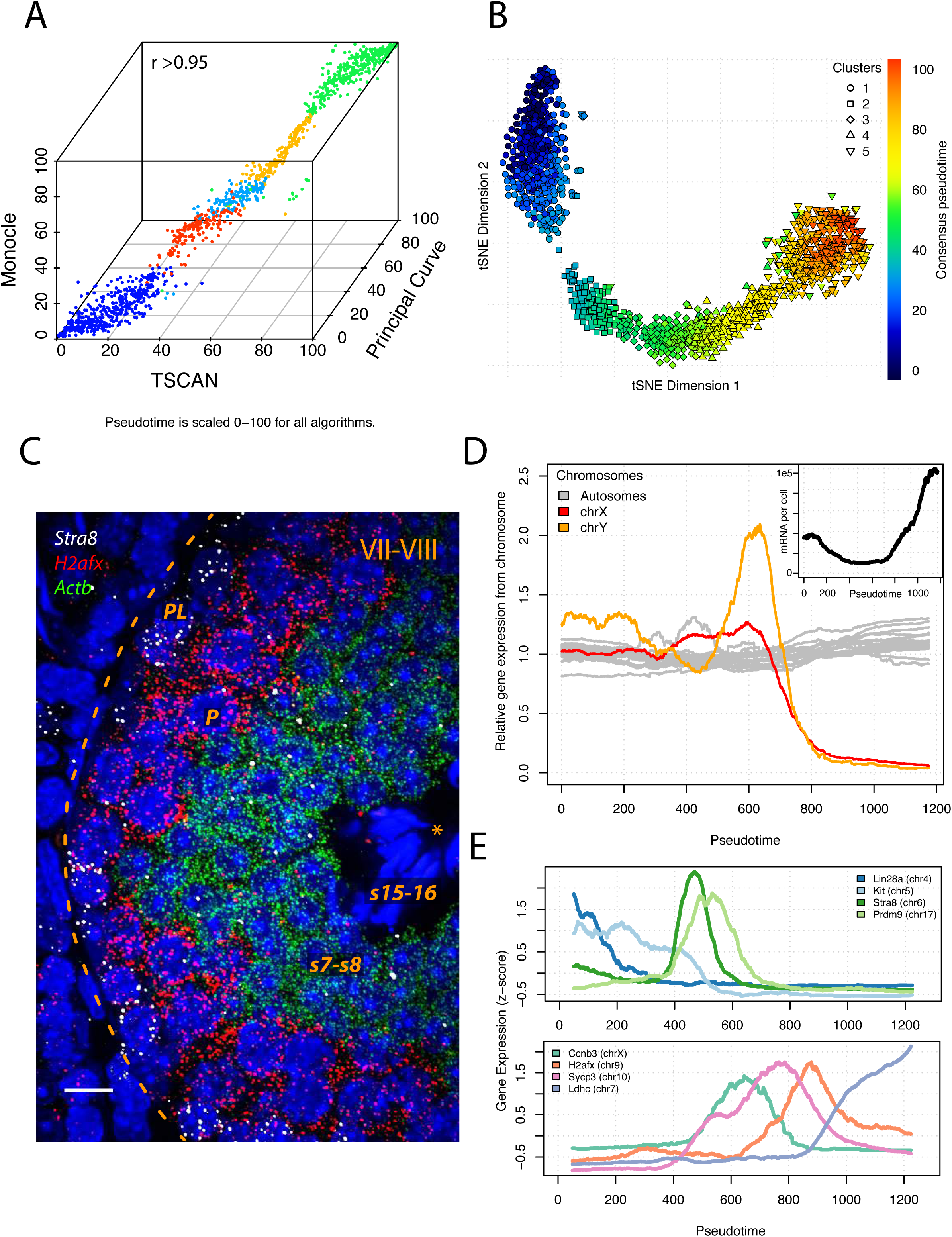
Multiplexed trajectory reconstruction is consistent with established expression patterns and allows to explore gene expression at high temporal resolution. a.) Pseudotemporal ordering is consistent across 3 different reconstruction methods (min. pairwise *r* > 0.95) and recapitulates the of stages (colors) defined earlier by marker genes. Each cell is a dot is a cell, ordered in 3 dimensions according to the pseudotime calculated by each algorithm (scaled 0-100) and colored as in Fig. 1. b.) Consensus trajectory (color) is the average of the 3 methods and is represented on the tSNE map. Clusters are denoted by symbols. c.) Three color single-molecule RNA FISH shows sequential expression of two consecutive prophase markers (*Stra8*, **Pl**: Preleptotene; *H2afx*, **P**: Pachytene) and the *Actb*, specifically upregulated in post-meiotic cells. Roman numeral: spermatogenic stage; **s1-16**: step 1-16 spermatids; * marks the lumen and dashed line marks the base of the tubule; scale bar: 10 μm. d.) Relative expression of autosomes (grey) and sex chromosomes (X: red, Y: orange) throughout prophase-I. While autosomes are steadily expressed, sex chromosomes are sharply downregulated before pachytene. Rolling average of normalized chromosomal read counts are scaled to by the median expression along the trajectory. The inset shows the raw transcript counts per cell as detected by sequencing, dropping before leptotene, and sharply increasing in pachytene. e.) Expression of later stage marker genes from (Fig. 1) and other key genes resolve expression dynamics at a higher resolution. Early and late genes are separated in the top and bottom panel for clarity. Gene expression is displayed as z-score normalized rolling averages (window: 100). Temporal reconstruction precisely predicts expression timing. *Lin28* peak is restricted to earliest *Dazl*+ spermatogonia, unlike *Kit*, and that the *Stra8* slightly precedes *Prdm9*, as reported.

The next two clusters (3 & 4) displayed the peak expression of *Dazl*, *Sycp1* & 3. They also a shared expression of zygotene specific genes involved in recombination, such as *Hormad1* (Shin *et al*., 2010) and *Meiob* (Luo *et al*., 2013), as well as *Elavl3*, *Gml* and *Mnd1* (Data S2). A key difference is that while cluster 3 shows high expression for many sex chromosomal genes (e.g. *Scml2*, *Fmr1*), cells in clusters 4 and 5 only express residual amounts of these mRNAs.

Taken together we concluded that clusters 1 to 5 show unique gene expression patterns, which recapitulated established stages of prophase-I. We built an online explorer for all genes (https://vertesy.shinyapps.io/ExpressionBrowser/) and listed all differentially expressed genes (Table S1-2). We also found subpopulations within these clusters, suggesting finer transitions. To analyze these systematically, we extended our computational analysis to a staging-free approach, better reflecting the continuity of spermatogenesis.

### 3. Multiplex trajectory reconstruction precisely recapitulates established meiotic events

Multiple recent approaches successfully reconstructed pseudotemporal trajectories from single-cell transcriptomes using either principal curves (Petropoulos *et al*., 2016), TSCAN (Ji and Ji, 2016), or Monocle 2 (Qiu *et al*., 2017). We reconstructed temporal order in our data set by all three methods (Fig. S3). It resulted in a similar, but not identical ordering (min. pairwise *r* > 0.95) (Fig. 2A, Online Methods). From the three reconstructions we calculated the consensus trajectory reflecting developmental time (Fig. 2B), which allowed us to analyze transcriptome dynamics in a more reliable and precise way than possible before.

First of all, this map recapitulated the previously determined stages, while revealing much finer events, as detailed below (Fig. 2E). We then used *in vivo* samples to validate this reconstruction, by 3-color single-molecule RNA fluorescence in situ hybridization (smFISH). We selected the (pre)leptotene gene *Stra8* (cluster 2) (Anderson *et al*., 2008), the pachytene specific *H2afx* (cluster 4) and the gene *Actb*, which is primarily expressed in spermatids (Ventela *et al*., 2000). The smFISH clearly pinpointed spatially-separated populations. Cells formed a sparse outer layer of *Stra8*+ (pre)leptotene cells decorating a dense intermediate layer of *H2afx*+ pachytene cells, enclosing an inner ring of *Actb*+ spermatid cells (Fig. 2C). Additionally, the three key transcriptional events (silencing at meiotic entry, meiotic sex chromosome inactivation and pachytene activation) are correctly recapitulated (Fig. 2D).

We then focused on finer differences. The reconstruction precisely predicted that *Lin28a* is expressed only in the earliest stage spermatogonia (Chakraborty *et al*., 2014). It also accurately predicted that *Stra8* peaked shortly before *Prdm9* (Sun *et al*., 2015). Remarkably, all other analyzed marker genes also peaked in the expected developmental order (Fig. 2E). Altogether, we concluded that the consensus trajectory faithfully recapitulated the developmental progress of early spermatogenesis, thus, it allows for a reliable, in-depth computational analysis of gene expression patterns.

### 4. Temporal expression maps show continuous differentiation from (pre)leptotene to pachytene

Next, we analyzed all genes’ expression profiles along the trajectory. First, we classified gene expression dynamics, and found that most genes showed a single expression peak (Fig. 3A). We then analyzed the co-expression profiles of genes peaking around (pre)leptotene. This showed how the meiotic master regulator *Stra8* expression preceded other preleptotene genes. As an example, we highlighted the cascade of *Stra8, Tex15* and *Dmc1* activation within one stage (Fig. 3B, selected by highest z-score).

**Fig. 3:**
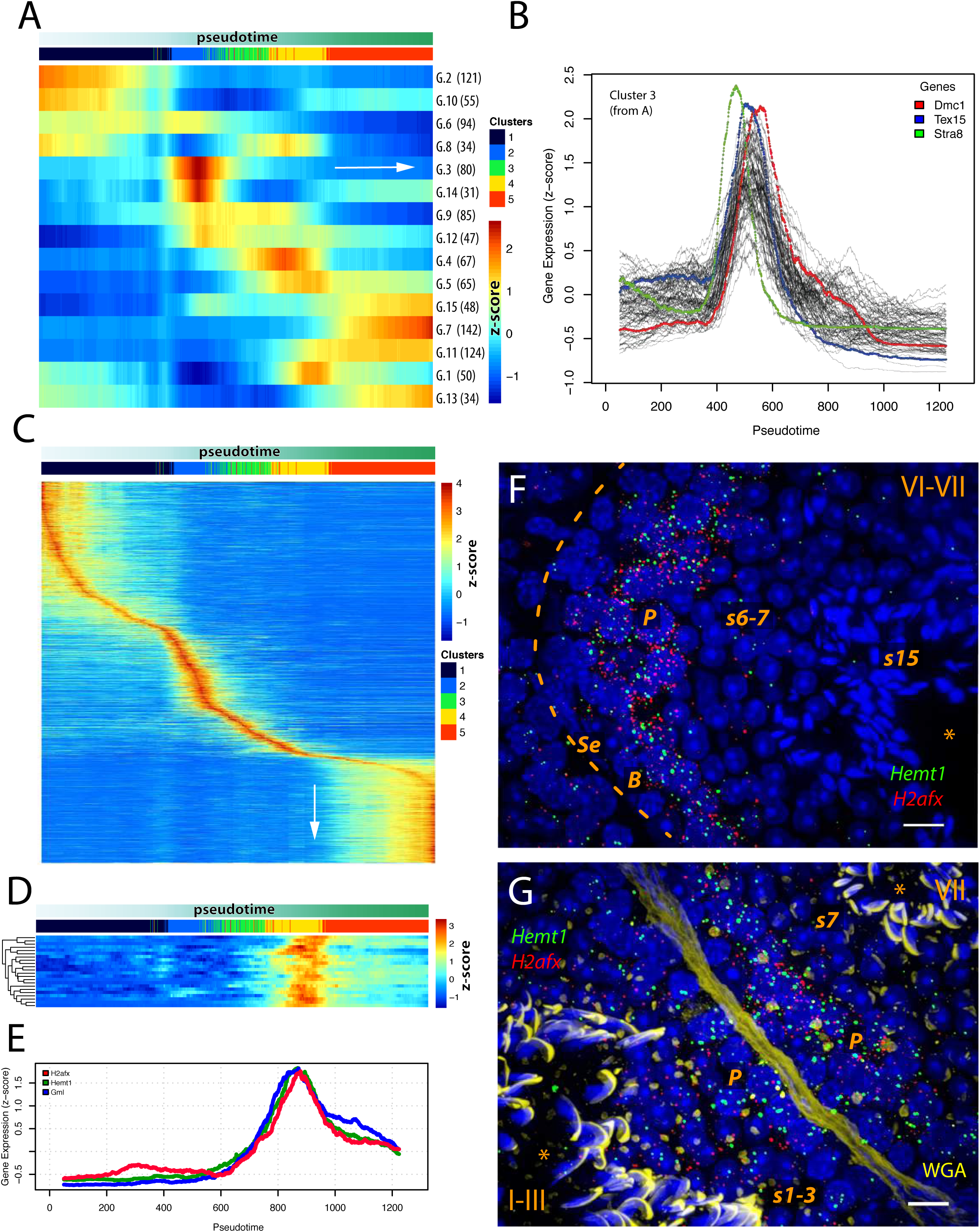
Systematic analysis of expression timing and co-expression patterns finds novel co-expression patterns. a.) K-means clustering of the 1077 highly expressed genes identifies specific peaking time for each gene. Clustering was done on z-transformed moving averages of normalized gene expression. b.) Traces of all genes in class 3 show a clear expression peak between 400 and 600. The 3 sharpest peaking genes (by maximum z-score) are highlighted in color. c.) Z-score normalized rolling average of raw transcript counts of all genes with a single, sharp peak, ordered by their peaking times. Note that the number of unique genes specific to the first (spermatogonia) and last (pachytene) cluster is inflated by nonspecific genes expressed beyond the analyzed time window. d.) Search for similar expression dynamics identified multiple genes (e.g: Ube2t, Hdac1, Trim28) co-expressed with *H2afx*. e.) *Gml* (*Hemt3*) vs *Gml2* (*Hemt1*) are the most stringently co-expressed genes with *H2afx*. f.) Single-molecule RNA FISH shows the co-expression of *Gml2* and *H2afx* in Pachytene (**P**) spermatocytes *in vivo*. Roman numeral: spermatogenic stage; **s1-16**: step 1-16 spermatids; * marks the lumen and dashed line marks the base of the tubule; scale bar: 10 μm. WGA staining omitted for clarity. g.) Single-molecule RNA FISH with wheat germ agglutinin staining (WGA, in yellow). WGA stains tubule boundaries and acrosomes, thus it allowed the staging of tubules in combination with nuclear morphology. Two adjacent tubules of different stages. The earliest pachytene cells are found in tubules of stage I-II, when pachytene cells just start to upregulate pachytene specific genes. Scale Bar: 10 μm

To understand this sequential activation deeper, we selected genes with a single expression maximum along the trajectory (Online Methods, Fig. S4). We then ordered these genes by time of peaking (Fig. 3C). First, it showed the three most unique stages in our dataset: spermatogonia, (pre)leptotene and pachytene. By showing the number of genes changing at each stage, it also highlighted the magnitude of changes upon meiotic entry, and pachytene activation. The analysis also suggests that sequential and sharply timed gene activation might be the regulatory clockwork of spermatogenesis, possibly triggering genes with wider expression profiles as seen in (Fig. 3A, B).

This analysis allowed us to identify genes co-expressed with known marker genes, which in turn can help to characterize their function. As an example, we identified 21 genes (r > 0.9) with expression patterns similar to *H2afx* (Fig. 3D). The best correlating genes were Hemt1 (Gml2) and *Gml* (*Hemt3*) (Fig. 3E), a pair of uncharacterized genes, although *Gml* was previously identified in prophase spermatocytes (Xue *et al*., 1999).

To validate this stunning co-expression of *Hemt1* and *H2afx*, we performed smFISH (Fig. 3F-G). This confirmed their co-expression within a spatially-restricted population, which suggests a co-functionality. To help researchers finding other co-expressed genes, we implemented an interactive co-expression browser as part of our online toolbox. Taken together, we created a compendium of gene expression dynamics in meiotic prophase-I and developed interactive tools for further investigation.

### 5. Silencing of an *n-Myc* transcription factor network coincides with global silencing during meiotic entry

Previous studies elegantly inferred transcription factor binding sites for large clusters of cells (Green *et al*., 2018) or clusters genes (Jung *et al*., 2018) during meiotic entry. Building upon this, we aimed at understanding the transcriptional programs behind the global silencing during meiotic entry. Therefore, we compared the transcriptome of spermatogonial and (pre)leptotene cells (Fig. 4A, Table S3). We analyzed these by identifying significantly enriched GO-terms of >4-fold differentially expressed genes (Online Methods, Table S3). The GO-terms were then clustered according to semantic similarity (Supek *et al*., 2011).

**Fig. 4:**
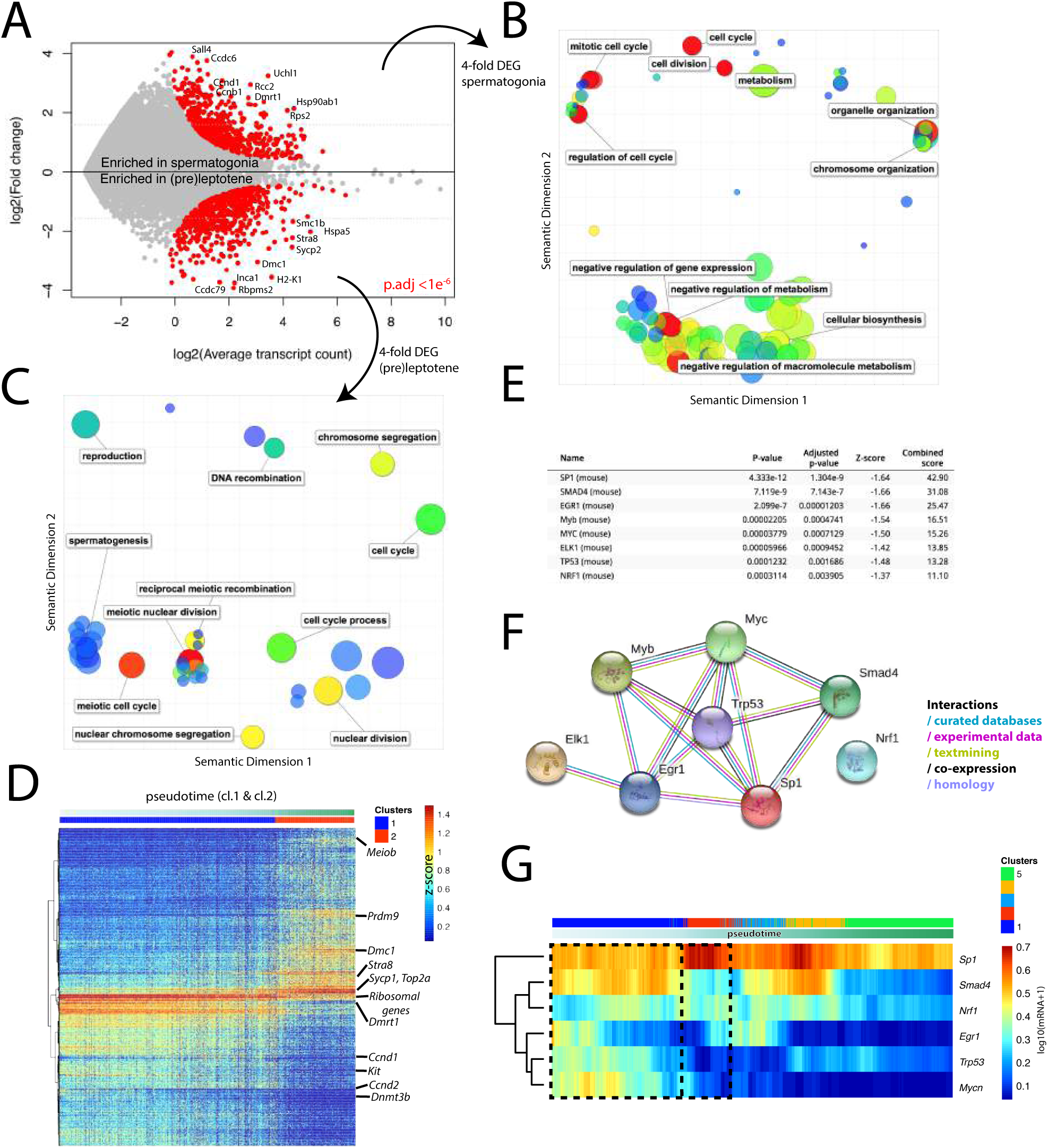
Meiotic entry is coupled to a downregulation of a tightly connected hub of transcription factors, and coincides with metabolic and transcriptional silencing. a) Pairwise differential gene expression between spermatogonia (cl.1) and (pre)leptotene (cl.2) cells shows 113, and 109 genes at least 4-fold enriched in either cluster, at p.adj<1e^−6^ (highlighted in red, Benjamini-Hochberg corrected). Selected gene names are displayed. All significant genes are listed in (Table.S4). b) Semantic similarity map shows key, stage specific biological processes. Significant GO-terms within the 4-fold spermatogonia (**B**), and (pre)leptotene (**C**) enriched genes-as identified in (A)-are grouped by similarity. Dot sizes correspond to the number of genes associated, colours denote the −log10(False Discovery Rate). Representative GO-terms with low FDR and high gene count are labeled. c) See above. d) Z-score normalized expression heatmap of at least 2-fold, significantly up- or downregulated genes in spermatogonia or (pre)leptotene cells. Cells in each cluster are ordered along the trajectory. Selected representative genes are labeled. e) List of 8 transcription factors with significantly enriched (p.adj. < 0.001) binding sites within spermatogonia specific 2-fold enriched genes, using the TRANSFAC and JASPAR databases. f) The identified transcription factors form a densely connected protein-protein interaction network as identified by STRING. g) Rolling average (window: 100) normalized gene expression of the detected transcription factors along the trajectory. Dashed lines indicate approximate spermatogonial and (pre)leptotene boundaries along the trajectory.

Besides the clear and expected switch from mitotic to meiotic program, the transcriptome of (pre)leptotene cells showed strong metabolic silencing (Fig. 4B, C). Metabolic silencing also may explain the drastic decrease of mRNA content in late spermatogonia, which lasts until pachytene (Fig. 2D). Many housekeeping genes were silenced, and their expression generally varied throughout the entire prophase-I (Data S3).

To better understand how meiotic entry is regulated, we focused on transcription factors. As many transcription factors are expressed at the detection limit of single-cell sequencing, they often cannot be reliably identified by differential gene expression analysis. Therefore, we analyzed the transcription factor binding site enrichment in the promoters of differentially expressed genes (Online Methods) (Matys, 2006; Portales-Casamar *et al*., 2010; Kuleshov *et al*., 2016). To gain statistical power, we analyzed >2-fold enriched genes in either cluster (Fig. 4D, enriched GO-terms recapitulated previous results, see Table S3).

Eight transcription factors showed highly significant (p.adj. <0.01) binding site enrichment in the spermatogonial genes (Fig. 4E). In contrast, no transcription factor was unique to the (pre)leptotene stage, while some (*Sp1* and *Smad4*) showed a highly significant enrichment in both gene sets (Table S3).

We found that six out of eight of the predicted transcription factors from a tightly interconnected hub (Fig. 4F, Online Methods) based on protein interaction network (Szklarczyk *et al*., 2017). This pinpoints a core regulatory network may regulate the mitotic-meiotic transition. When testing these predictions on our data, we found that most predicted spermatogonia-specific factors (*n-Myc, Trp53, Egr1, Smad4*) are silenced expectedly. Curiously *Sp1* –which cooperates with *Smad4* in TGF-β signaling– showed the opposite trend (Feng, Lin and Derynck, 2000). *Nrf1* (Wang *et al*., 2017) was expressed continuously, while *Myb* and *Elk1* were not detected (Fig. 4G).

Taken together, explicit silencing-rather than activation-drives the transcriptional switch in meiotic entry. The silencing showed a particularly strong metabolic control. *Myc* is known to regulate more than 11% of all genes, in particular metabolic ones (Fernandez, 2003). Hence, the downregulation of the spermatogonia-specific *n-Myc* variant (Braydich-Stolle et al., 2007) may explain the global transcriptional silencing upon meiotic entry.

### 6. 34% of the sex chromosome genes are induced right before sequential inactivation by MSCI

The second major wave of transcriptional silencing in prophase-I is the meiotic sex chromosome inactivation (MSCI). This leads to the global silencing of the sex chromosomes (Henderson, 1963; McKee and Handel, 1993). While post-meiotic silencing has been analyzed in detail (Ernst *et al*., 2019), and multiple recent single-cell studies referred to MSCI (Chen *et al*., 2018; Jung *et al*., 2018; Lukassen *et al*., 2018). However, the patterns of gene silencing during MSCI, and the underlying regulation remained open questions.

To tackle this question, we classified all sex chromosomal genes along the trajectory. This confirmed their silencing at late zygotene, but also showed considerable gene-to-gene variability in silencing dynamics (Fig. 5A). Four sets of these genes formed distinct waves of transcription. The largest wave contains genes expressed in spermatogonia (cluster 1).This, in turn, is consistent with the observations that the X chromosome is enriched in premeiotic genes (Khil *et al*., 2004; Koslowski *et al*., 2006). Yet, interestingly, many other genes were transiently induced prior to MSCI. The combination of these patterns (gradual silencing & pre-MSCI transcriptional burst) also clarifies why the total expression from both the sex chromosomes ramps up before MSCI (Fig. 5B). We also confirmed our observation using a recently published dataset (Ernst *et al*., 2019).

**Fig. 5:**
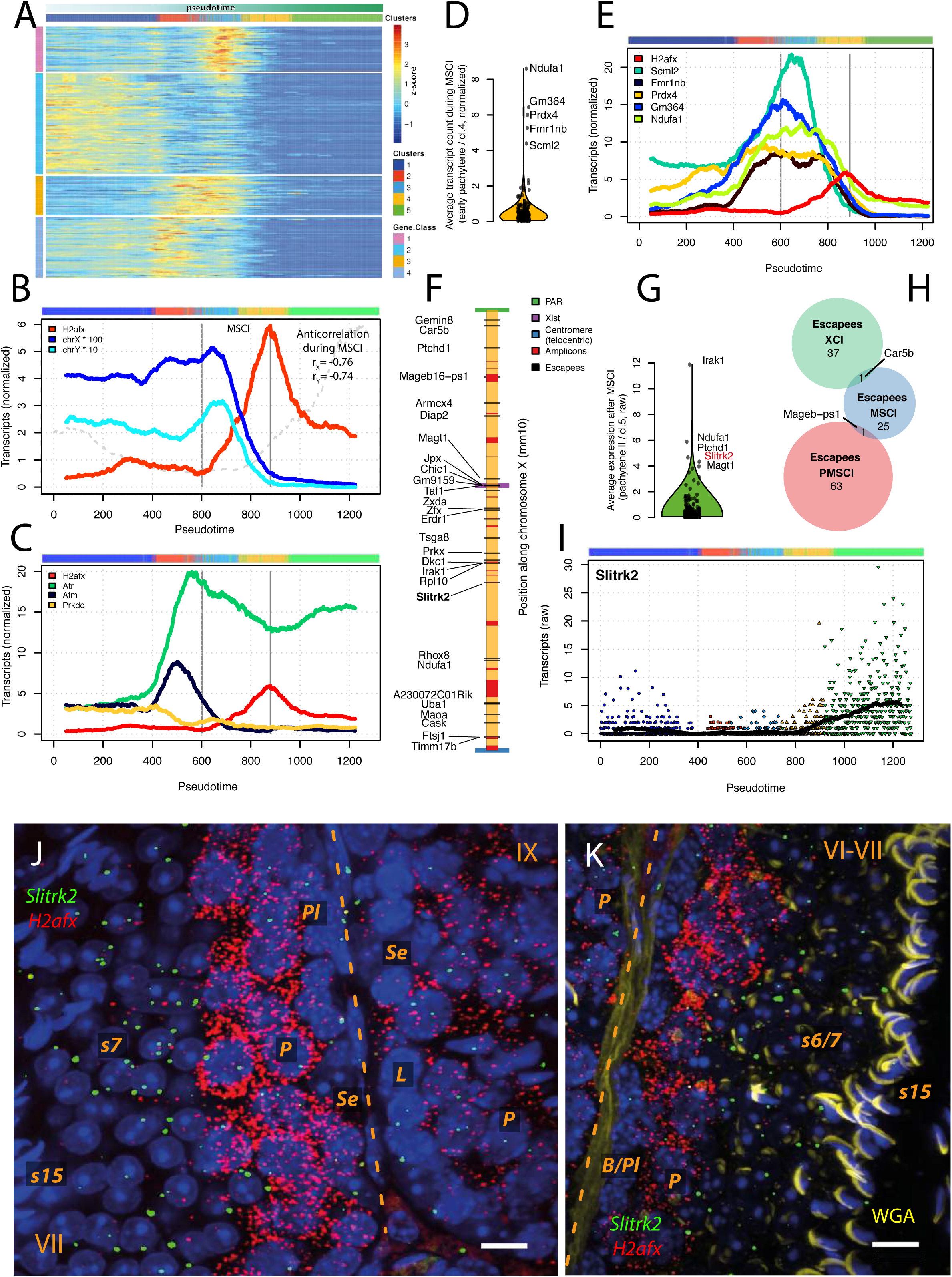
A wave of sex chromosomal gene activation precedes meiotic sex chromosome inactivation (MSCI) a.) 34% of sex chromosomal genes are induced specifically before and silenced during MSCI. Genes are classified into 4 classes by k-means clustering of Z-score normalized rolling average gene expression profiles. Class 1 and 3 contains genes induced before and silenced by MSCI. Class 2 contains spermatogonia specific genes, while class 4 contains leptotene specific genes, and genes which are steadily expressed until MSCI. b.) Both sex chromosomes show increasing relative expression before a rapid silencing that coincides with the induction of *h2afx* gene. Rolling average normalized expression values are divided by 100 for X, or by 10 for the Y chromosome for visualization. Dashed lines indicate the start and maximum of *h2afx* transcription, also marking the start and end of MSCI. Pearson correlation coefficients between the sex chromosomes’ and *H2afx*’s expression during MSCI are denoted (X, Y, respectively). Grey dashed line indicates transcript counts per cell, as in Fig.2D. c.) Gene expression of known post-transcriptional regulators of *H2afx*. As reported *Prkdc* is not expressed and *Atm* is only expressed earlier, during double strand break formation. The sole activator expressed during MSCI is *Atr*, and its’ expression is unchanged after (pre)leptotene, suggesting that MSCI may not only be controlled at the post-transcriptional level. d.) Average expression of all peaking genes (class 1&3) genes during MSCI (cl. 4) shows that a small set of genes is very highly expressed when meiotic silencing is already in progress. e.) Individual traces of the highly expressed peaking genes (from **D**) shows a late and sharp silencing dynamics. *H2afx* expression (red), and MSCI boundaries (grey vertical lines) are displayed as a reference. f.) Chromosomal location of the 27 sex chromosomal genes (all from chr. X) with an average expression above 1 in pachytene (cl.5). g.) Highly expressed escapee genes. Average expression of sex chromosomal genes in pachytene II (cl.5) identifies a handful of escapee genes that are highly expressed after MSCI. h.) The ampliconic gene *Mageb16* is the only MSCI-escapee gene, which is also described to escape post-meiotic sex chromosome inactivation (PMSCI), and *Car5b* is the only gene that is also described as an escapee in female X chromosome inactivation. i.) *Slitrk2* is the only gene whose expression anti-correlated with the total expression from the X-chromosome. While it is efficiently silenced before, it starts to be expressed during MSCI and increases throughout pachytene. j.) Single-molecule RNA FISH shows that *Slitrk2* escapes during late pachytene (right, stage IX) and it maintains post-meiotic expression (left) spermatids *in vivo*. Roman numeral: spermatogenic stage; **Se**: Sertoli cell; **L**: Leptotene; **P**: Pachytene; **s1-16**: step 1-16 spermatids; dashed line marks the base of the tubule; scale bar: 10 μm. WGA staining omitted for clarity. k.) Single-molecule RNA FISH with WGA staining (in yellow) shows continued *Slitrk2* expression in spermatids, but it is not expressed in the earliest pachytene cells (cl. 4). Scale Bar: 10 μm.

We also noticed that the sex chromosomal expression clearly anti-correlated with the expression of *H2afx* (Fig. 5B, stages marked by a color bar on top). This observation contrasts with the ubiquitous deposition of *H2afx* along the genome, and with the idea that is mostly regulated post-transcriptionally (Turner *et al*., 2004; Bellani *et al*., 2005; Turinetto and Giachino, 2015). *H2afx* can be phosphorylated by *Atm, Atr* and *Pkrdc*, and we confirmed that only *Atr* is expressed during MSCI (Bellani *et al*., 2005), however it shows decreasing mRNA levels (Fig. 5C). The lack *Atr* induction around MSCI alongside the observation that *H2afx* is induced precisely at this time suggests that the timing of MSCI is partly controlled by *H2afx* transcription. Possibly, then the main role of the *Atr* could be to activate the newly deposited histones.

It has been proposed that MSCI is simply a selective lack of reactivation of the sex chromosomes during pachytene (Page et al. 2012). Here, we show that MSCI is an separate silencing event (Fig. 5B). First of all, sex chromosomal transcription sharply falls before pachytene activation. Second, many sex chromosomal genes were specifically upregulated before silenced (Fig. 5A). Finally, transcript levels after (pre)leptotene silencing are stable until pachytene, except from the sex chromosomes (Fig. 2D).

Some X chromosomal genes have autosomal paralogs, which switch expression to the autosomal copy upon MSCI. We indeed see the timing and extent of autosomal compensation in our data. As examples, we detected that when the X chromosomal gene *Rpl10* is silenced by MSCI, its paralog, *Rpl10L* on chr12 is induced (Jiang *et al*., 2017). The paralog pairs *Pgk1* (chrX) – *Pgk2* (chr17), and the *Cetn2* (chrX) – *Cetn1* (chr18) (Hart *et al*., 1999) show the same trend (Data S6).

It is unknown whether MSCI follows any pattern and if genes are silenced in any particular order. The best-studied example of chromosome-wide silencing is the X chromosome inactivation (XCI) in female mice. Although XCI evolved to achieve dosage compensation, both involve the silencing of the whole X chromosome. Silencing in XCI is primarily governed by chromosome structure and proximity to the X-inactivation center (Nora *et al*., 2012; Engreitz *et al*., 2013).

We therefore tested the possibility of a silencing center, or a silencing bias towards either chromosomal end. In the telocentric chromosomes of mice, centromeres occupy one end of each chromosome. For the X and Y chromosomes, the other end forms the pseudoautosomal region (PAR). While the centromere holds the sister chromatids together, the PAR holds the X & Y chromosomes together in meiosis. Thus, both ends might bias chromatin silencing.

We then analyzed the silencing dynamics along the X chromosome, on a linear basis. While we found ample variability in silencing dynamics, we detected no spatial effect towards either end of the X chromosome, nor towards the *Xist* locus (Data S4). Thus, we concluded that unlike X-inactivation, MSCI is unlikely to be governed by proximity to an inactivation center.

Genes in the pseudoautosomal regions (PAR) are X/Y homologous, and not silenced, so they could be used as an internal negative control for MSCI. In mice however, PAR only contains 3 active genes (Kasahara et al., 2010). While *Sts*, and *Asmt* (*Hiomt*) transcripts were not detected, *Mid1* (*Fxy*) indeed showed low, but constant raw transcript counts throughout prophase-I (Fig. S6E).

We further examined the expression of the latest silenced genes in the second half of MSCI (cluster 4). We found a small set of genes with remarkably high expression (Fig. 5D). These genes –except *Prdx4*– were specifically induced in zygotene and showed high expression, when the expression of other sex chromosomal genes was already decreasing (Fig. 5E). *Scml2* is a testis-specific polycomb protein, which regulates the ubiquitination state of *H2afx*. It showed one of the strongest and latest expression peak, underlining its important role in controlling MSCI (Luo *et al*., 2015) and genes escaping silencing (Adams et al. 2018).

Based on the functional cues among the late-silenced genes, we performed a comprehensive analysis of gene function during silencing. We classified sex chromosomal genes into fast- and slow-silenced categories based on their characteristic silencing time (Online Methods). For each group, we calculated GO-term enrichment as compared to all sex chromosomal genes (Online Methods). We found that while early-silenced genes were typically localized to the nucleus (43% of the genes, FDR < 4e^−3^). Inversely, late-enriched in genes encoding mitochondrial proteins (19%, FDR < 2e^−5^). This further supported the notion that silencing is a coordinated process starting with regulatory genes and followed by cytoplasmic and metabolic genes (catalytic activity: 34% of late genes, FDR < 8e^−3^) (Table S4).

### 7. *Slitrk2* and other genes escape MSCI

We also noticed that the *Ndufa1* (Fig. 5D), is not only the last gene to be downregulated, but it also maintains a steady absolute transcript level throughout pachytene, possibly by escaping inactivation. Whether genes escape MSCI is debated in recent literature (Chen *et al*., 2018; Jung *et al*., 2018). Transcripts detected in pachytene can either be due to mRNA preservation over several days, or due to an active transcription from MSCI. Therefore, it is important to characterize if such genes show increased expression after MSCI, and to validate their expression pattern independently.

To examine if other sex chromosomal genes are also detected after MSCI, we calculated the average raw transcript counts in cluster 5 (as normalized counts may be misleading, see Fig. 2D and Online Methods). We identified 27 genes on chromosome X with above and an average of one transcript per cell (Fig. 5G). This correspond to approximately 10-100 transcripts per cell, given the estimated sensitivity of single-cell methods (Grün, Kester and van Oudenaarden, 2014). Importantly, we detected more transcripts for these 27 genes after MSCI, suggesting that their expression is not simply due to mRNA preservation.

*Slitrk2* showed the most striking expression pattern, as it was specifically upregulated after MSCI (Fig. 5G). *Slitrk2* is the only gene expressed from the hybrid sterility locus (*Hstx1/2*). It has been described to escape silencing in *domesticus X musculus* hybrids (Bhattacharyya *et al*., 2014). It is not expressed in Sertoli cells (Zimmermann *et al*., 2015), and we validated its expression *in vivo* by smFISH (Fig. 5J-K).

Mapping these 27 escapee genes on the X chromosome showed a clustered pattern around the *Zfx*, *Irak1*, and *Xist* loci (Fig. 5H). Note, that *Xist* itself is not expressed in males. ATAC-seq has previously identified the *Xist* locus as one of the most open chromatin regions during XCI in females (Giorgetti *et al*., 2016), which may contribute to the lack of silencing in genes located in proximity of *Xist*. In addition, we noticed that many of the escaping genes from the *Xist* locus are long, and often non-coding genes. Indeed, the transcript length of the 27 escapee genes are significantly longer than other X chromosomal genes (one-sided MWW test, p<0.006).

When compared to known escapees in post-meiotic sex chromosome inactivation (Namekawa *et al*., 2006) and MSCI, only *Mageb.ps-1* is shared. Similarly, when compared 38 XCI escapees (Berletch *et al*., 2015) with MSCI, only *Car5b* was common (Fig. 5I). These observations suggest that a mechanism of silencing and escaping distinct from previously studied cases (e.g. XCI) occurs during the meiotic phase.

Finally, we wanted to know if escapee genes show specific expression dynamics during prophase. Therefore, we clustered gene expression profiles along the trajectory. We found that most genes follow a pattern similar to *Ndufa1*, while the expression pattern of *Slitrk2* is unique (Fig. S5A). Yet, both patterns appeared as residual rather than functional. This is because relative transcript counts –within the expanding pachytene transcriptome– remained in all cases low. Escape from silencing then might simply be the consequence of open chromatin amidst global transcription. An exception to this might be Slitrk2.

Taken together, we found that a third of sex chromosomal genes are specifically induced before silencing by MSCI. Although MSCI is a short event in meiosis, we found remarkable differences in the silencing time of genes. While broad chromosome structure does not appear to affect silencing time, we found functional cues. early-silenced genes are mostly nuclear, but late-silenced genes are often cytoplasmic or mitochondrial. This leads to the interesting possibility of a “regulation first, processing second” silencing pattern. Yet, to understand the role of MSCI, one must consider a combination of ancestral and newly acquired functions (Turner *et al*., 2004). MSCI might have evolved for one reason, *e.g.* global silencing of unsynapsed chromatin, and once in place, it could evolve to new regulatory functions.

### 8. Genes in pachytene are activated in the functional order of mature sperm anatomy

The transient silencing during leptotene and zygotene ends before pachytene (Monesi, 1965). This leads up to 9-times more cellular mRNA than in earlier stages (Fig. 2D, inset, at 900 on the trajectory). In accord with a global transcriptional activation, cells in pachytene reach the most diverse transcriptome in our dataset, as quantified by entropy (Fig. S4E).

To characterize this massive transcriptional wave, we first identified genes upregulated in pachytene by differential gene expression analysis (Fig. 6A). We selected all genes that showed a 3-fold upregulation compared to all other cells (Online Methods). We classified these into 3 categories by k-means clustering of z-score normalized rolling average expression profiles (Fig. 6D). Clusters showed reproducibility >98% over 10000 bootstrapping rounds, and gene classes displayed sharp, distinct activation profiles (Fig. 6B & E).

**Fig. 6:**
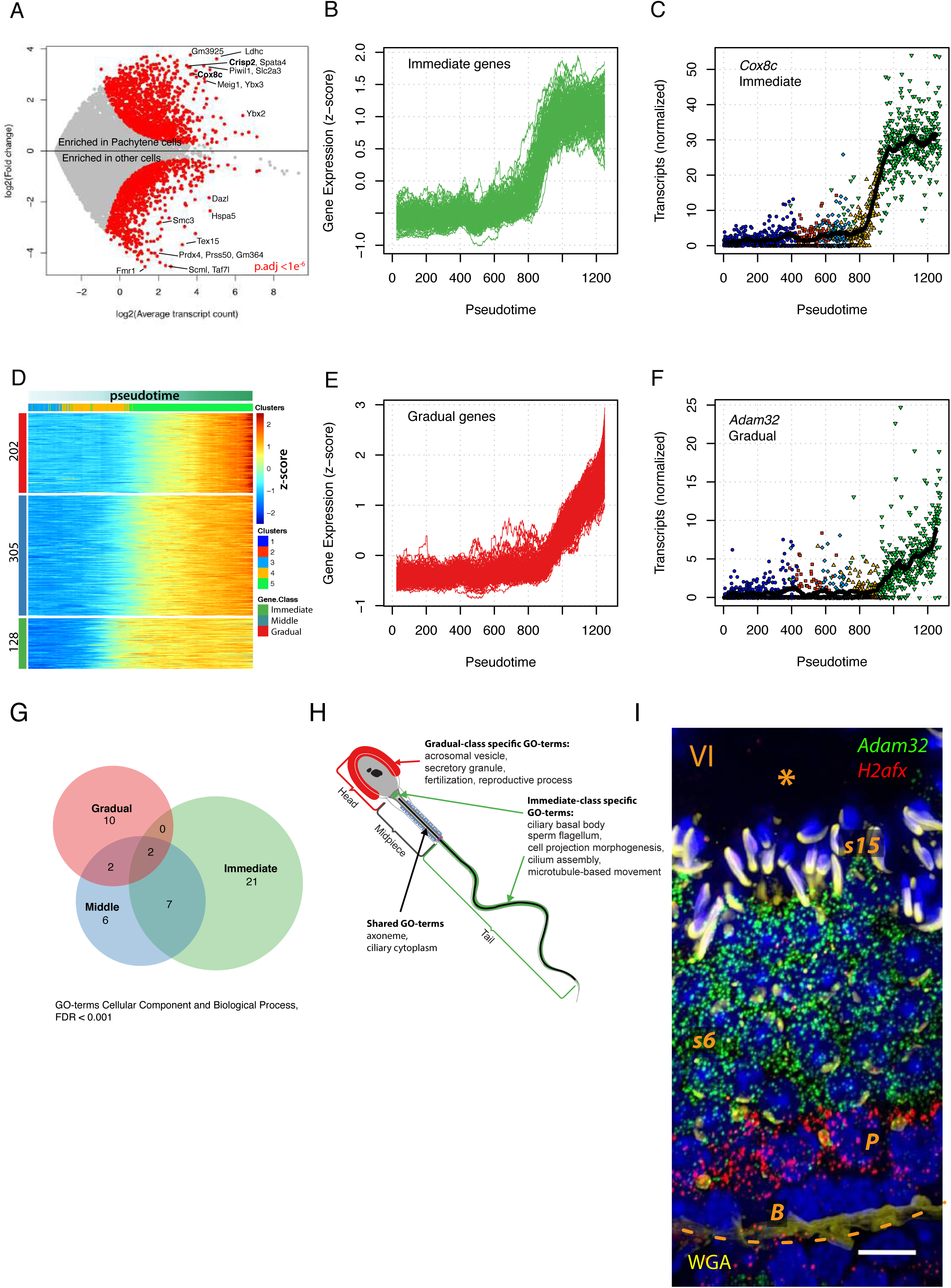
Genes in pachytene are induced in the same order as they are utilized in the mature sperm. a.) Differential gene expression analysis identifies 635 genes that are at least 3-fold upregulated in pachytene cells as compared to all other cells (p.adj < 1e^−6^, in red, Benjamini-Hochberg corrected). Representative genes are labeled in black. b.) Individual expression profiles of immediate genes show the step-like up regulation after MSCI. c.) The nuclear encoded mitochondrial gene, Cytochrome C Oxidase Subunit 8C (*Cox8c*) is prime example of sharply upregulated genes. d.) Gene expression profiles along the trajectory of the 635 genes were classified into 3 categories according to the shape and slope of increasing gene expression. K-means clustering was performed on z-transformed moving average gene expression. e.) Gradual genes all showed a slowly increasing gene expression in pachytene. f.) The uncharacterized *Adam32* gene is suggested to be involved in sperm-egg fusion and it shows the typical gradual accumulation of fertilization related genes. g.) Significantly enriched GO-terms suggest markedly different function between the immediate and gradual gene-sets (GO Biological Process and GO Cellular Component, FDR<1e^−3^). Immediate genes are enriched in energy and mobility related functions and structures, while the gradually upregulated genes are enriched in fertilization related functions and structures. h.) Gradual- and immediate-class specific GO-terms map to different anatomical regions of the sperm. Both unique and shared enriched GO-terms are analyzed, and informative GO-terms are displayed. Gradual-class specific terms and corresponding anatomical structures are in red. Immediate-class specific terms and structures are in green. Shared terms and the related axoneme is indicated by the black arrow. The full list of GO-terms is accessible in (Table S6). i.) Single-molecule RNA FISH shows the upregulation of the fertility related *Adam32* after pachytene and its post-meiotic expression in spermatids *in vivo*. Roman numeral: spermatogenic stage; **B**: B spermatogonia; **P**: Pachytene spermatocyte; **s1-16**: step 1-16 spermatids; * marks the lumen and dashed line marks the base of the tubule; WGA staining for acrosomes is in yellow; scale bar: 10 μm.

Many immediate genes had functions linked to the pachytene transcriptional activation itself. Other immediate genes encoded proteins for energy production (e.g. The mitochondrial cytochrome c oxidase genes) (Fig. 6C). Piwi-pathway related genes formed the next wave in the ‘intermediate class of genes (*Piwil1*, *Tdrd1*, *Pld6* and *Gtsf1*), protecting against transposons (Aravin, Hannon and Brennecke, 2007).

Yet, most genes activated in pachytene have post-meiotic roles (Da Cruz, 2016). Many of the enriched functions are first relevant in elongating spermatid (e.g. flagella formation) or in mature spermatozoa (e.g. binding to zona pellucida). These genes are thus surprisingly transcribed more than a week before their utilization. The *Adam* gene family (Primakoff and Myles, 2000) is a typical example of the pachytene-activated, fertilization-related genes. One of these gradual genes is *Adam32* (Fig. 6F). We validated its expression by smFISH (Fig. 6I), which also matched the expression pattern found in neonatal testis (Choi *et al*., 2003).

We also found distinct functional signature of immediate and gradual genes, while the intermediate class of genes showed a blend of functional enrichments (Fig. 6G). Interestingly, the genes were activated in parallel order to the order they will be used in the mature spermatozoa. While immediate genes play a role in flagella formation and cell movement, late genes are involved in fertilization (Table. S6). Accordingly, enriched GO-terms in the immediate and gradual classes mapped to different anatomical regions of the spermatozoa (Fig. 6H).

Surprisingly, while these genes are transcribed in a precise order, they are only translated and used more than a week later. This temporal discrepancy highlights the important role of mRNA storage in spermatogenesis. In accord, we noted that pachytene activation is exactly the time when *Dazl*, an RNA binding protein, is downregulated (Fig. 1D). Altogether, we observed a sequential pattern of gene activation in pachytene cells that reflect the spatial anatomy of the mature sperm. This temporal pattern might show a spatial pattern of maturation of the germ cell, from the tail to the head.

## Discussion

Currently, meiotic entry in mammals is not well understood, as highlighted by our inability to induce meiosis purely *in vitro*, without the co-culture of explanted tissues. To gain a transcriptome-wide and time-resolved insight, multiple recent studies applied single-cell mRNA sequencing combined with pseudo-temporal reconstruction (Chen *et al*., 2018; Green *et al*., 2018; Hermann *et al*., 2018; Jung *et al*., 2018; Lukassen *et al*., 2018; Ernst *et al*., 2019). These studies provided excellent resources and greatly contributed to understand the whole span of spermatogenesis. Yet, additional questions require deeper, fine-tuned, computational analysis.

Our study focused on early meiosis up until pachytene. We found that in contrast with the multitude of morphological stages, there are only three major transcriptomic events. In the first major transition spermatogonia committed to meiosis. This is accompanied by a global shrinkage of the transcriptome and a clear functional switch. Cells shift from a mitotic and catabolic to a meiotic and recombination-related transcriptome. This functional switch however appears to be regulated by a one-sided silencing of a transcription factors hub around *n-Myc*, while no comparable network is upregulated.

The second major transition is meiotic sex chromosome inactivation. It is thought that the phosphorylation of the silencing histone variant *H2afx* controls the timing of inactivation. We however found that *H2afx* gene is induced exactly during MSCI and it is anti-correlated to sex chromosomal expression. These together suggest that the timing of MSCI is controlled by the surplus of *H2afx* expression. At the same time *Atr* expression is flat, which may hint towards no temporal control on the post transcriptional level.

We then characterized chromosome-wide activation and silencing patterns before, and during MSCI. We found that sex chromosomal genes are silenced at distinct times, challenging established models for MSCI. For example, if repression would only appear to protect DNA, we would not expect that genes silence in a coordinated, sequential manner.

In general, silencing follows two major patterns: gradual silencing and transient activation followed by silencing. We found that 34% of sex chromosomal genes were specifically induced before MSCI. We speculate that some of these gene products may serve as buffer or pool until autosomal genes take over their functions in pachytene. Additionally, many of these transiently activated genes were involved in regulating gene expression. Interestingly, the chromatin regulator *Scml2* (Hasegawa *et al*., 2015; Luo *et al*., 2015; Adams *et al*., 2018) is one of the genes strongest induced before, and latest silenced upon MSCI, highlighting that gene function and silencing time are concerted. In addition, genes activated just before MSCI might act as a trigger for the third major transition, the pachytene activation.

Taken together, sex chromosomes are silenced in a functional, rather than structural order, raising the possibility that MSCI is involved in gene regulation. This, in turn, could also explain why MSCI is essential for meiotic program. Nevertheless, we think there is no single purpose behind MSCI: gene regulation could have evolved atop of its ancestral functions. To deeper understand MSCI, one could complement chromatin profiling with single-cell mRNA-seq, as for PMSCI (Ernst *et al*., 2019), or perform single-cell chromatin accessibility profiling.

Finally, the third major transition in prophase-I is pachytene activation. This global transcriptional activation boosts cellular mRNA content up to 9-fold. We found that immediately and gradually activated genes are functionally distinct. While immediate genes were related to energy metabolism and spermatozoal movement, gradual genes were related to fertilization. Accordingly, the corresponding gene products mapped exclusively to the posterior, or anterior parts of the mature sperm.

In sum, we elucidated key events in of early meiosis, and provided a transcriptome-wide, precisely timed, gene expression atlas for online exploration. Our approach hopefully will be followed by functional studies showing the importance of timing in meiosis.

## Acknowledgements

We would like to thank Mauro help for experimental help, Willy Baarends and Joost Gribnau for valuable comments on the manuscript, and help with histological analysis, Stefan van der Elst and Reinier van der Linden for help with FACS sorting, Nicolas Battich for advices on microscopy analysis, Josi Peterson-Maduro for logistic help and Fanny Sage for the mouse drawing.

## Supplementary figure legends

**Fig. S1. Filtering and quality control for clustering**

(**A**) Transcript count per cell shows a bimodal distribution. Low quality cells below 3000 transcripts are discarded (red). This results in 1274 high quality cells (green) with a median of 20488 reads.

(**B**) Saturation determines the optimal cluster number in RaceID2. With increasing number of clusters, the Within Cluster Dispersion (WDC) decreases. When further increase of the number of clusters (**k**) yields a decrease (grey) that is within the estimated error interval (red) of the decrease, **k** is selected (5 in this case).

(**C**) Gap Statistic did not yield a peak in single cell mRNA-seq datasets, therefore cannot be used to unambiguously determine the cluster number. Of not that it also saturates around 5, the cluster number determined by the saturation of within cluster dispersion.

(**D**) The silhouette plots showed the average similarity of each cell to all cells in its own cluster, relative to other cells in the closest neighboring clusters. Positive values for most cells suggest that they are well represented within their own cluster.

(**E**) High Jaccard index shows the high reproducibility of clusters across multiple bootstrapping rounds. The Jaccard index is calculated as the intersect over the union of cells in the same cluster across multiple bootstrapping rounds, pairwise. RaceID qualifies clusters reproducible above Jaccard index 60% (or 0.6).

(**F**) Sequencing depth does not explain observed transcript count differences. Average over-sequencing per cell varies across individual cells a magnitude more than across clusters. Average oversequencing per cluster is lower in clusters with high transcript counts (cl1 & 5) as expected if the high read counts were explained by biology, and not by sequencing depth.

**Fig. S2. Differences in transcriptome and clustering results are not explained by technical effects.**

(**A**) Distribution of cells across cell types, which were identified by k-medoids clustering.

(**B-D**) Cells from all four libraries (**B**), all three animals (**C**), and cells prepared with either CEL-Seq1 or CEL-Seq2 primers (**D**) -- denoted by color, respectively -- contribute equally to all cell types (denoted by shapes), as shown on the t-SNE map.

(**E-F**) Detected Gene and Transcript counts per cell detected both transiently decrease from meiotic entry (cl.1→ cl.2) until a global transcriptional activation in pachytene (cl.5).

**Fig. S3. Pseudotemporal ordering by three different algorithms**

(**A**) Principal graph of cluster centers, and cell projections as reconstructed by Monocle 2 using DDRTree algorithm. Colors denote cell clusters as determined by RaceID, colored by Monocle2.

(**B**) Minimum spanning tree of cluster centers, as reconstructed by TSCAN (in black). Cells were subsequently projected on this tree. Colors denote cell clusters identified by TSCAN using a multivariate Gaussian mixture model in PCA space.

(**C**) Reconstructed Principal Curve in t-SNE space and minimal cell projections, to the curve. Colors denote cell clusters as determined by RaceID. Clustering results are not used in this reconstruction, only provided for easier comparison.

(**D**) Pairwise scatterplots and Pearson correlation coefficients across the 3 reconstruction methods.

**Fig. S4. Filtering Peaking Genes by Quantile Ratio**

(**A**) The ratio of the 100% and 95% quantiles of each gene’s expression efficiently distinguishes peaking genes (high maximal z-score) caused either by outlier expression values, or by representative gene expression. While these genes both have high z-score maxima (31 and 17 respectively), quantile ratio of two example genes clearly separates outlier genes like *Prm1*. Log quantile ratio shown by dotted lines for *Zfy2* (left) and *Prm1* (right).

(**B-C**) Normalized transcript counts in single cells and rolling averages (in black, window: 50) of *Prm1* and *Zfy2*. Maximum z-scores of rolling averages are high in both cases, however in the case of *Prm1*, this is clearly caused by one outlier cell with 1000 transcripts, thus it does not reflect a general change in expression.

(**D**) Entropy quantifies the diversity of the transcriptome per cell and showed that pachytene-II cells (cl.5) had transcriptome diversity even higher than spermatogonial stem cells.

**Fig. S5. Expression patterns during and after MSCI in cluster 4 & 5**

(**A**) Correlation network of gene expression plotted in PCA space. Spearman correlation was calculated on the normalized expression patterns of sex chromosomal genes along the trajectory within cluster 4 & 5. The blue arches represent high Spearman correlation coefficients. All parameters were left as default in the *corrr* R-library. Most genes are like *Ndufa1* (big cluster on the left), in that their normalized expression (relative to the total transcript count per cell) decreases, while absolute transcript count might stay at a similar level. Sporadically, escapee genes show particular expression patterns, such as genes similar to *Ptchd1*, showing flat relative, increasing absolute counts, or a strong increase such as *Slitrk2*.

(**B**) The double peaking expression pattern of the multicopy *Rbmy* gene family.

**Fig. S6. Changes in molecular chaperones’ expression coincides with major transcriptional changes, (activation and silencing).**

(**A**) Constitutively expressed (housekeeping) *Hsp90ab1* (90α-class b) and *Hspa8* expression decreases upon meiotic entry, alongside with 3 chaperones involved in protein import to mitochondria (*Hspd1*, *Hspe1*, and *Hspa4*)

(**B**) From (pre)leptotene to zygotene a mixed set of chaperones is induced. The constitutive *Hsp90b1* shows moderate induction (∼20-30 mRNA/cell), while the endoplasmic reticulum related, inducible *Hspa5* shows very strong upregulation (∼70-80 mRNA/cell). The signaling related small HSP member *Hspb11* shows a minor (∼5mRNA/cell), but distinct peak at meiotic entry.

(**C**) The induced *Hsp90α* is known to be involved in cell cycle regulation, and it is strongly induced upon meiotic entry, reaching ∼150 transcripts per cell. Beside its role in stress response, *Hsp90α* is also induced in response to the induction of excessive translation. At the same time moderate to low induction of small HSPs *Hspb6* and *Hspa4l*, and the *Hsp70* family member Hspa2, which is suggested to be involved in spindle integrity.

(D) Log10 scaled moving average gene expression of all displayed chaperones. Importantly, immediate early genes (as defined in van den Brink et al 2017) are mostly lowly expressed in the dataset, or specifically expressed at different stages (see also at Data S5).

(**E**) Expression of *Mid1 (Fxy)*, the only detected genes from the pseudoautosomal region (PAR).

## Supplementary table descriptions

**Table S1: Differential gene expression: each cluster vs. all others**

Differential gene expression analysis of each cluster as compared to all other clusters. Summary for each cluster (number of genes enriched), summary for each gene (arithmetic and geometric mean expression per cluster) and the enrichment statistic per gene, per comparison are provided.

**Table S2: Differential gene expression: stepwise comparisons**

Differential gene expression analysis of each cluster compared to the previous (in developmental order) cluster. Summary for each cluster (number of genes enriched), summary for each gene (arithmetic and geometric mean expression per cluster) and the enrichment statistic per gene, per comparison are provided.

**Table S3: Changes of gene expression and function during meiotic entry.**

TRANSFAC and JASPAR enrichment of transcription factors, binding to all at least 2-fold enriched genes; enrichment statistic per gene, per comparison; and the list of significantly enriched GO-terms in gene products of cluster 1 and 2 cells are provided.

**Table S4: Changes of gene expression and function during meiotic sex chromosome inactivation**

Characteristic silencing time and peak gene expression (rolling average) of sex chromosomal genes during MSCI; List of early-silenced genes and respective significantly enriched GO-terms; the same information for late-silenced genes; and finally, the list of sex chromosomal genes, which were used as a statistical background for the enrichment analysis.

**Table S5: Changes of gene expression and function during pachytene activation**

Cluster assignment of upregulated genes (k-means, k=3); list of and description of gene products for each cluster; and respective lists of enriched GO-terms are provided.

## Supplementary dataset descriptions

**Data S1 Transcriptome wide clustering as well as marker gene expression supports gradual, as opposed to switch-like differentiation of spermatogonia**

A. *Dazl*+ spermatogonial transcriptome does not sub-cluster along *Kit* or *Lin28a*/*Sox13*/*Plzf* expression boundaries. Spearman or Pearson correlation based clustering of spermatogonia confirms overall transcriptional similarity observed in Fig. 1 regardless of the expression of Kit or the markers of undifferentiated spermatogonia. The cells that cluster apart are of lower quality, as defined by raw transcript counts.

B. Expression of reported markers of *Kit*-, undifferentiated spermatogonia. All spermatogonial cells exclusively express *Dmrt1*. 77% of spermatogonia express *Kit* (≥1 raw transcript), and only these cells also express Cyclins (*Ccnd1, Ccna2, Ccng1*). *Kit*-*Dazl*+ (and *Dmrt1*+) spermatogonia exclusively express a set of markers for undifferentiated spermatogonia: *Plzf* (or *Zbtb16*), *Sall4*, and *Sox13*. However, the expression of other markers for undifferentiated spermatogonia, such as: *Uchl1*, *Sox3*, *Stella* (*Dpp3*), or *Lin28a*), are not restricted to the Kit-population.

**Data S3 Differential Gene Expression**

Visual summary of differential gene expression analysis (Table S1-S2). Panels 1-5: comparison of cells in each cluster with all other cells. Panels 6-9: comparison of each cluster with the developmentally earlier cluster. Genes in red are significantly differentially expressed at p <1e^−6^ (Benjamini-Hochberg corrected).

**Data S5 Stress.Genes.van.den.Brink.2017**

Dissociation-induced genes (covering most early immediate genes) as defined in (Van Den Brink *et al*., 2017), are mostly lowly expressed. Expressed “dissociation-induced genes” show varied dynamics along prophase-I and do not point to population of cells, which is particularly affected by the isolation procedure.

## Supplementary online methods Vertesy et al

### Mice strains, Organ Isolation and Dispersion

To separate germ cells from somatic cells, as well as to selectively sort germ cells from meiotic prophase-I, we used a mouse line expressing Dazl-GFP-V5 on a bacterial artificial chromosome (BAC). In this construct, the GFP sequence is integrated directly before the stop codon, and its expression is controlled by Dazl promoter and enhancer sequences (Chen *et al*., 2014). *All mice were housed according to the Hubrecht Institute guidelines, which include access to food and water ad libitum. All animal experiments were carried out in compliance with standards for care and use of laboratory animals, with approval from the Dutch Animal Experiment Committee of The Royal Netherlands Academy of Arts and Sciences (KNAW). Mice were between 2 and 3 months old at time of sacrifice*.

In the first step, we isolated testicles from Dazl-GFP mice and chopped them into pieces. *Next, these tissue pieces were transferred to a 50 mL falcon tube and incubated in a collagenase type II (2 mg/mL Gibco, 17101015)/phosphate-buffered saline (PBS, Lonza, 17-516F) solution placed in a water bath set at 37 °C for 1 hour to degrade the extracellular matrix and release cells from their niche. We used 10 mL collagenase solution per TA*. After this initial incubation, the tissue pieces were incubated in Trypsin at 37 degrees with gentle intermittent shaking for 8-10 min until the islets were dispersed into single cells. The dispersed tissue was washed briefly with cold PBS then filtered through a 25 μm sieve to remove debris and aggregates.

### FACS Sorting

The Dazl-GFP mice allows us to select germ cells from spermatogonial up to pachytene spermatocyte stages, as Dazl expression in adult testes is restricted to germ cells of these stages. We used Propidium Iodide (PI) to exclude dead cells and VybrantViolet (<u>cat. V35003</u>) staining for DNA content to exclude haploid cells.

We incubated the cells with PI and VybrantViolet on ice before FACS-sorting, according to manufacturer’s protocol. Cells were kept on ice until sorting. We used FACSJazz (BD biosciences) with the following configuration for all sorts Channel, Dye, Function: [488] 530/40, GFP, Dazl expression; [488] 585/29, PI, Live cells (PI negative); [405] 450/50, VybrantViolet, DNA content. Channel interpretation: [Excitation wavelength] Emission wavelength / Detection bandwidth.

First, we selected single, live cells based on PI exclusion and scattering (P1 and P2 and P3). Next, we used GFP expression and VybrantViolet staining of Live Dazl-GFP positive single-cells. Finally, the selected cells were sorted into hard shell 384-well plates (Bio Rad, cat HSP3801) containing 200 nl of RT primers, dNTPs and synthetic mRNA Spike-Ins. To avoid evaporation, 5 μl of vapor-lock (QIAGEN, cat 981611) was added on top.

After sorting, we spun down and froze the plates immediately. The plates were stored at −80°C until processing. For details see (Van Den Brink *et al*., 2017). For library preparation and sequencing we followed the SORT-seq protocol (Muraro *et al*., 2016).

Sorting experiments were done in biological triplicate and technical replicates. There were no relevant differences between replicates that would affect the conclusions. We both used CEL-seq1 and CEL-seq2 based primers. They showed differential sensitivity for a set of genes, which we effect we normalized out. See ***Analysis*** for consequences and normalization.

### Single-Cell mRNA library preparation

We prepared cells into sequencing libraries by SORT-seq. For the detailed protocol, see: (Muraro *et al*., 2016) and (Van Den Brink *et al*., 2017). In brief, we used a 3-prime, poly-A based, UMI-corrected library construction protocol. SORT-seq is an improved and robotized version of the CEL-seq2 protocol designed for FACS sorted cells (Hashimshony *et al*., 2016).

### Microscopy, single-molecule RNA fluorescence in situ hybridization (smFISH)

For microscopy, testis isolated from 2-3 months old male wild type mouse were fixed in 4% RNAse free PFA at room temperature for 4 hours and cryopreserved by incubating overnight in 30% glucose solution at 4C. Tissue was then cryomounted on OCT, frozen and sectioned into 10μm thick slices. Tissue sections were then used for performing single-molecule RNA-FISH using the ViewRNA ISH Cell assay kit (Invitrogen: QVC0001). Tissue sections undergo a new fixation with 4% RNAse free PFA for 20 min at RT, permeabilization and protease treatment steps. Next, gene specific probe sets are hybridized (∼20 oligos per gene), followed by sequential branched-DNA (bDNA) signal amplification, and finally individual mRNA molecules are visualized. The probes were designed to hybridize with the mRNAs of Stra8 (VB6-16903-VC), H2afx (VB1-3030165-VC), Actb (VB4-10432-VC), Gml2 (VB6-3223290-VC), Slirtk2 (VB6-3218124-VC), Adam32 (VB6-3221253-VC). Nuclei were stained by Hoechst, lectins were stained by Wheat Germ Agglutinin (WGA, molecular probes #W11261). Sections were imaged using a 63x objective on a PerkinElmer Ultraview VoX spinning disk microscope combined with a Leica SP8. Images were analyzed in ImageJ/FIJI.

### Histology and staging of seminiferous tubules

Seminiferous tubules a classified into 12 stages (I-XII) based the presence of different cell types (Russell *et al*., 1993). Many cell types are indistinguishable by themselves under the microscope but can be inferred if the stage of the tubule is known, because only certain cell types are present at each tubule stage. Staging of tubules is based on nuclear and acrosomal morphology, as well as the position and presence of certain cells types (Meistrich and Hess, 2013). We staged tubules by fluorescence microscopy using a combination of fluorescent lectin conjugates and nuclear morphology (Nakata *et al*., 2015). Stages are indicated by Roman numerals. Each image was recorded in 4 fluorescent channels, either DAPI + 3 smFISH probes, or DAPI + WGA + 2 smFISH probes. For clarity, DAPI + smFISH images are shown in the main figures, while WGA staining is not always shown.

### Data Analysis

#### Alignment and Quantification

Paired-end reads from Illumina sequencing were aligned to the mouse RefSeq transcriptome with BWA-MEM (Li, 2013). We trimmed low quality bases and discarded low quality and ambiguous alignments. For details on alignment, see (Grün *et al*., 2015). After alignment, we removed UMI-duplicates from uniquely mapping reads. UMI-duplicates are reads from the same cell, same gene sharing the same unique molecular identifier revealing that they are were amplified from the same original mRNA molecule. Next, based on binomial statistics, we converted the number of observed UMI-s into transcript counts (Grün, Kester and van Oudenaarden, 2014).

#### Filtering

First, we selected cells with >2000 transcripts, yielding 1274 high quality cells. Next, we selected genes with at least 2 transcripts in at least 20 (out of 1274) cells. These cells had on average a median of 8079 transcripts from 1077 highly expressed genes.

#### Downsampling and normalization

Samples were normalized by taking the average of 10 downsampling rounds. Generally speaking, downsampling is better at keeping the statistical properties of the data, such as noise level, while normalization has no loss of information and therefore can be more sensitive. We combined these two approaches for an optimum solution. For downstream analysis, including k-medoids clustering, we used RaceID (Grün *et al*., 2016) and custom scripts.

#### Normalization of libraries generated by CEL-seq1 and CEL-seq2 primers

As the SORT-seq protocol was designed during the time course of this study, we prepared libraries with two types of primers for reverse transcription. The original version is described in (Grün, Kester and van Oudenaarden, 2014), and it was based on CEL-seq1 (cs1) (Hashimshony *et al*., 2012). Later, an optimized primer version was developed (Muraro *et al*., 2016), which was based on the CEL-seq2 (cs2) protocol (Hashimshony *et al*., 2016). We prepared 490 high quality (filtered) cells by cs1-, and 766 by cs2-primers.

To normalize the 2 protocols, first we calculated the average expression per gene over all index sorted plates per cs1 and cs2 batches. Next, we calculated the ratio of the batch averages, and normalized all genes to have the same average expression in both batches. After normalization we filtered highly expressed genes which are then used for clustering. Genes which had a more than 10-fold bias before normalization or were 4 times more often detected in either protocols were excluded from clustering. In Fig. S2 we show that cells from each batch contribute equally to all clusters regardless of primer design, biological replicate, or batch of library preparation.

#### Clustering and tSNE

For many steps of analysis, including k-medoids clustering of cells and tSNE projection, we used an in-house improved version of RaceID2 (Grün *et al*., 2016). We implemented parallel computing in RaceID2 and extended its functionality to handle pseudotime related analysis and better accommodate multi-panel plotting. The code will be freely available on https://github.com/vertesy/RaceID-v2.5-Multithreaded upon acceptance, with restriction to attribute the original author of RaceID upon use.

#### Establishing the number of clusters for k-medoids clustering

In RaceID, the optimal cluster number is determined as the saturation point of the Within Cluster Dispersion (WDC). By separating the data into more (thus smaller) clusters WDC decreases. When adding a new cluster yields a decrease that is no bigger than the estimated error interval of the decrease, k is selected. In this estimation, we repeatedly clustered cells into 1 to 10 preset clusters, and calculated WDC for each number of clusters. The optimal cluster number was estimated to be 5.

For comprehensiveness, we also calculated gap statistics (Tibshirani, Walther and Hastie, 2001). Gap statistic in single cell mRNA-seq datasets often does not yield a peak, therefore cannot determine the cluster number unambiguously. Nevertheless, it also saturates around 5, as estimated before

The silhouette plot shows the average similarity of each cell to all cells in its own cluster relative to all cells in the closest neighboring clusters. Positive values for most cells suggest that they are well represented within their own cluster. High Jaccard index shows high reproducibility of clusters across multiple bootstrapping rounds. The Jaccard index is calculated as the intersect over the union of cells in the same cluster across two bootstrapping rounds. RaceID qualifies clusters reproducible above Jaccard index 60%, while Jaccard indices in this study range 74-99%.

#### Differential Gene Expression

For differential gene expression analysis, we used RaceID2. The algorithm was specifically designed to model transcript count variability in single-cell data by a negative binomial noise model (Grün, Kester and van Oudenaarden, 2014), building on (Anders and Huber, 2010). P-values of significantly up or downregulated genes were calculated based on Poissonian statistics and corrected for multiple testing by the Benjamini-Hochberg method (Grün and van Oudenaarden, 2015).

We found that fold change as commonly calculated as the fraction of arithmetic mean (“average”) gene expression in compared cell types is not well suited for single cell data, as suggested previously (Bengtsson *et al*., 2005).

Analysis based on arithmetic mean often identifies genes, which we consider false positives when looking at the raw expression data, in single cells. This is because arithmetic mean is a metric sensitive to outliers. It often indicates a big differences, which are not representative of the group of cells; in fact they are regularly caused by a few outlier cells with very high expression.

Median is commonly used as an outlier robust metric of the central tendency, it however loses sensitivity at low integer read counts, which is the typical range the for majority of the genes in single cell data. Furthermore, median becomes meaningless if near 50% of the cells have 0 expression (also known as zero-inflated data). Subtype specific genes with a narrow expression window are often overlooked by using median (e.g.: Zfy1 & 2).

Geometric mean is both robust against outliers and handles zero-inflated data, and it was found to be the appropriate metric for gene expression data that varies on the log-scale (Vandesompele *et al*., 2002; Bengtsson *et al*., 2005; Goymer, 2005). We therefore used the fold change of geometric means alongside the classical definition of fold change of arithmetic means, so that a gene must satisfy the same threshold for both metrics. This largely decreased our false positive detection, while maintaining similar sensitivity.

#### Expression patterns defining cluster boundaries

Groups of cells can be defined by either a unique expression of genes or by a combination of multiple non-unique expression patterns. We compared gene expression in each cluster to all other clusters to find uniquely expressed genes (TableS2), and we also compared gene expression in each cluster to the previous cluster (=developmental stage) to find gradual or stepwise changing genes (Table.S3).

### GO-term enrichment

We found that GO-term enrichment on coding genes gives cleaner results, therefore we calculated GO-term enrichments using the STRING (v10.5) protein interaction database (Mering, 2003) everywhere, unless stated otherwise.

#### Functional enrichment and REViGO Analysis for spermatogonial and (pre)leptotene genes

We first performed pairwise differential gene expression between spermatogonial and (pre)leptotene clusters, then calculated GO-term enrichment on 4-fold enriched genes in either clusters. Next we analyzed the significantly enriched GO-terms (Biological Process, FDR < 0.05) using REViGO (Supek *et al*., 2011), with setting (similarity=0.9, semantic similarity measure: SimRel, associated values: −log10(FDR), GO term sizes: Mus musculus). REViGO first removes redundant (nested) GO terms, then calculates a 2 dimensional multidimensional scaling map based on semantic similarity of the terms. The full list of GO-terms is provided in (Table S3).

#### GO annotation defined gene sets

We first identified relevant GO-terms using EBI’s QuickGo (Binns *et al*., 2009) within the GO-term hierarchy. Next, we downloaded mouse gene annotations for each term, excluding negative annotation qualifiers using the AmiGO browser (Carbon *et al*., 2009). We selected genes detected in our dataset and analyzed genes as described per analysis.

#### Pseudotemporal ordering of single cells

To draw a comprehensive picture of all genes changing expression at different times, we needed to reconstruct their developmental trajectory (Cannoodt, Saelens and Saeys, 2016). As we did not measure time, but infer, how long cells progressed in spermatogenesis, this procedure is called *pseudotemporal ordering*. We ordered cells according to gradual, transcriptome wide changes in gene expression by 3, distincts and established methods: Monocle 2 (Qiu *et al*., 2017), TSCAN (Ji and Ji, 2016) and principal curve (Hastie and Stuetzle, 1989) as used in (Petropoulos *et al*., 2016). The 3 methods are compared in their main features, dimensionality reduction, clustering, and trajectory modeling below.

Table: Comparison of the trajectory inference methods used.

| Approach | Dimensionality Reduction | Clustering | Trajectory model | Ref |
| --- | --- | --- | --- | --- |
| Monocle 2 | DDRTree (discriminative dimensionality reduction via learning a tree) | k-medoids, provided by RaceID2 | Principal Graph (a Principal Curve with branches allowed) learnt during dimensionality reduction by DDRTree. | (Qiu *et al.*, 2017) |
| **TSCAN** | PCA (Principal Component Analysis) | Multivariate gaussian mixture in PCA space | Minimum spanning tree of cluster centers | (Ji and Ji, 2016) |
| **Principal Curve** | tSNE (*t*-distributed stochastic neighbor embedding) | None | Principal Curve | (Hastie and Stuetzle, 1989) |

#### Monocle 2

Monocle 2 identifies differentially expressed genes across clusters of cells (provided from RaceID2 in this case), then uses Discriminative Dimensionality Reduction with Trees (DDRTree) (Mao *et al*., 2016), a Reversed Graph Embedding (RGE) algorithm to reconstruct a minimum spanning tree connecting cluster centroids, which is called principal tree. Cells then are projected to this principal tree, and their position along the backbone of the tree is counted as pseudo time. Monocle 2 uses an unsupervised inference method, where one does not need to feed the number of trajectories or paths, nor branching points. In fact multiple minor (<8% of the cells) side branches are detected by the algorithm, which all fall close to the main tree (Fig. S3).

#### TSCAN

In TSCAN, we provide the filtered normalized dataset. TSCAN uses PCA to reduce the dimensionality, which is followed by clustering by fitting a mixture of multivariate normal distributions using the Mclust package. The number of clusters here is determined by Bayesian information criterion, in our case it was 5. Next, TSCAN connects cluster centers with a minimum spanning tree, and finally it projects single cells on the tree, which is the cell order (Fig. S3 B).

#### Principal Curve

We fitted a principal curve (Hastie and Stuetzle, 1989), as in (Petropoulos *et al*., 2016) to the two-dimensional t-SNE map (Fig. 1 A) of the cells in 100 iterations. The curve is initialized as a line segment; we use first principal component line. Next, each point of the curve is recalculated as a local average of all data points that project to this points using a spline smoothing with 4 degrees of freedom (smooth.spline function). In the next round new projections are calculated, etc. Already after 10 iterations the curve converged. The final curve was calculated after 100 iterations (Fig. S3 C).

#### Comparison and Consensus or Multiplex Pseudotime

Projection coordinates in all 3 approaches are scaled to [0,100] and pairwise Pearson correlation coefficients are calculated: TSCAN vs. Principal Curve: 0.956; TSCAN vs. Monocle: 0.973; Principal Curve vs. TSCAN: 0.956. Essentially all 3 algorithms captured the linear progress from spermatogonia to pachytene spermatocytes (Fig. 2 C). To further increase robustness, we calculated the consensus, or multiplex pseudotime as the average pseudotime for each cell in all three trajectories calculated by the different algorithms //Table.S3.

#### co-expression analysis

We calculated the Pearson correlation coefficient (r) over the normalized gene expression of pseudotime ordered cells.

#### Gml (Hemt3) and Gml2 (Hemt1)

*Gml* (*Hemt3*) and *Gml2* (*Hemt1*) is an interesting pair of genes, which are encoded on opposite strands, ∼250 bases apart (ensembl mouse, GRCm38.p6, Chromosome 15: 74,806,933-74,812,856). While only Gml, but not Gml2 protein was shown to be expressed in spermatocytes in prophase (Xue *et al*., 1999), the two transcripts show 89% sequence identity, as determined by blast alignment of the cDNAs. Sequencing reads show roughly equal and high read counts for both genes, suggesting that this readout is not due to sequencing errors (in such a case, one transcript would have 1-10% of the reads from the more abundant transcript).

Both transcripts are extremely well correlated with *H2afx* along pseudotime (r1 r2 =??), and accordingly, the Gml signal is correlated with *H2afx* signal in smFISH. However, because of the sequence similarity, it is not possible to resolve *Gml* and *Gml2* by smFISH (transcripts are tiled along with 20+ oligos), therefore smFISH gives a readout of these transcripts together.

#### Early- and late-silenced genes on the sex chromosomes

First, we selected all sex chromosomal genes that had a running median expression maximum above 1 during MSCI (pseudo time points 650-900) as defined by the *h2afx* expression ramp (Fig. 5B). To quantify the pace of gene expression changes, we defined the characteristic timescale of silencing (TS) for each gene. TS is defined in a way similarly to a substance’s half-life: it is the time (pseudo time points) required for gene expression to drop to half of its maximal value, and it was corrected with an offset if the maxima did not coincide with beginning of MSCI. Without this correction, late activated genes, e.g. *Scml2*, would be mistakenly counted as an early-silenced gene. TS was calculated on rolling average normalized gene expression with a window of 100. Finally, we ordered all genes according to their characteristic silencing timescale and divided the distribution into earlier and later half. We note that some reads mapped to a microRNA, Mir684-1. It is a multicopy gene across autosomes and the X chromosome. For this reason, and since CEL-seq is not suited to study microRNAs, we considered it a mapping artifact, therefore we excluded it from the analysis.

GO-term enrichment was calculated on the interaction network of coding genes as described above. The statistical background for the enrichment were all coding genes on sex chromosomes present in the gene model used (mm10), (Grün, Kester and van Oudenaarden, 2014).

#### Escapees

As pachytene cells had ∼9 fold more reads than zygotene, relative expression levels might not be indicative of leaky silencing, therefore we calculated expression of unnormalized transcript counts. The variation among single cells was counteracted by calculating a rolling average over the cells ordered along the trajectory.

#### Classification gene expression patterns along the trajectory

First, the normalized gene expression values of individual cells were ordered along the trajectory. Next, we calculated the rolling average expression with a window of 50 (or 100 when indicated). To make genes with widely different expression levels comparable regarding their activation or silencing time, we z-score transformed these values. Finally, this gene expression matrix was classified by k-mean clustering.

#### Transcriptome wide identification of time-specific or peak genes

As we extended our analysis to lowly expressed, we found many lowly expressed genes with specific expression times, such as *Zfy2* (Fig. S3 C). Expression peaks in ordered series can be identified by maximum z-score, however single outliers cause artifactual high z-scores (Fig. S3 B), but in this expression regime, one cannot filter on expression level. Therefore, we calculated the quantile-ratio, a metric that can robustly remove artifactual peaks. In this, we calculate the ratio of the 100% and 95% quantiles of each gene’s expression. Genes only highly expressed in a few outlier cells have extremely high quantile ratio, and this metric efficiently distinguishes these genes from genes with true expression peaks (Fig. S3 A).

#### Analysis of Transcription Factor Binding sites

We found that meiotic entry from the spermatogonia towards preleptotene is characterized by global transcriptional silencing accompanied by the upregulation of defined set of genes. To find out which transcription factors driving these changes, we analyzed transcription factor binding sites enriched in the all genes that are at least 2-fold differentially expressed in cluster 2 vs 1, and vice versa. We used the online too *Enrichr* (Chen *et al*., 2013; Kuleshov *et al*., 2016), which searches across the TRANSFAC and JASPAR transcription factor binding profiles (Matys, 2006; Portales-Casamar *et al*., 2010). *Enrichr* scans the −2000 and +500 vicinity of transcription start sites for transcription factor binding sites. The list of transcription factors with significantly enriched binding sites is then analyzed in the 2017 version of the amazing STRING database to identify functional and physical interactions, as well as GO-term enrichment (Mering, 2003; Szklarczyk *et al*., 2017). We found that all upregulated genes are regulated by a tightly interconnected set of transcription factors.

### Data And Software Availability

All analysis was performed in R-studio v1.0.136. For single cell analysis we used the StemID algorithm (Grün *et al*., 2016) and custom scripts. Figures were generated by R packages pheatmap v1.0.8 (Raivo, 2013) and MarkdownReports v2.9.5 (Vertesy, 2017). Upon acceptance, the source code for analysis will be available “as-is” under GNU GPLv3 at https://github.com/vertesy/Spermatogenesis.Single-Cell. Sequencing data is deposited under accession number <u>GSE114788</u> at GEO.

